# Differential gene expression is associated with degeneration of mating-type chromosomes in the absence of sexual antagonism

**DOI:** 10.1101/626291

**Authors:** Wen-Juan Ma, Fantin Carpentier, Tatiana Giraud, Michael Hood

## Abstract

In animals and plants, differential expression of genes on sex chromosomes is widespread and it is usually considered to result from sexually antagonistic selection; however differential expression can also be caused by asymmetrical sequence degeneration in non-recombining sex chromosomes, which has been very little studied. The anther-smut fungus *Microbotryum lychnidis-dioicae* is ideal to investigate the extent to which differential gene expression is associated with sequence degeneration because: 1) separate haploid cultures of opposite mating types help identify differential expression, 2) its mating-type chromosomes display multiple evolutionary strata reflecting successive events of gene linkage to the mating-type loci, and 3) antagonistic selection is unlikely between isogamous haploid mating types. We therefore tested the hypothesis that differential gene expression between mating types resulted from sequence degeneration. We found that genes showing differential expression between haploid mating types were enriched only on the oldest evolutionary strata of the mating-type chromosomes and were associated with multiple signatures of sequence degeneration. We found that differential expression between mating types was associated with elevated differences between alleles in non-synonymous substitution rates, indels and premature stop codons, transposable element insertions, and altered intron and GC content. Our findings strongly suggest that degenerative mutations are important in the evolution of differential expression in non-recombining regions. Our results are relevant for a broad range of taxa where mating compatibility or sex is determined by genes located in large regions of recombination suppression, showing that differential expression should not be taken as necessarily arising from antagonistic selection.

**Author Summary:** Differences between males and females, from morphology to behavior and physiology, are considered to largely reflect differential expression of genes that maximize fitness benefits relative to costs that are specific to one sex. However, there is an unexplored alternative to such ‘sexually antagonistic selection’ to explain differential expression. Reproductive compatibility is often determined by genes located in large non-recombining chromosomal regions, where degenerative mutations are expected to accumulate and may separately affect the expression of alternate alleles of genes. We tested the role of genetic degeneration in determining differential expression between the isogamous haploid mating types of the anther-smut fungus, *Microbotryum lychnidis-dioicae*, where sexually antagonistic selection is not a confounding factor. We show that differentially expressed genes are highly enriched in the non-recombining mating-type chromosomes, and that they are associated with various forms of degenerative mutations, some of which indicate that the less expressed allele suffers greater mutational effects. Our finding of the role for degenerative mutations in the evolution of differential expression is relevant for a broad range of organisms where reproductive compatibility or sex is determined by genes in regions of suppressed recombination, and shows that differential expression should not be taken as necessarily arising from antagonistic selection.

## Introduction

Sexual antagonism occurs when trait values that increase gene transmission through the male function decrease gene transmission through the female function, or conversely [1–4]. Differential gene expression between sexes is widely thought to be the primary means of resolving the conflict of sexual antagonism, resulting in commonly called “sex-biased genes” [5,6]. This adaptive explanation for the existence of sex-biased genes predominates the literature [5–10], although empirical tests are rare and alternative hypotheses may explain the existence of sex-biased genes in sex chromosomes. Non-recombining regions on sex chromosomes often undergo degenerative changes that may also cause differential gene expression. Recombination suppression indeed renders selection less effective due to reduced effective population size [11], genetic hitchhiking of deleterious mutations with beneficial ones [2], and deleterious mutation sheltering [12–14]. Over time, deleterious mutations accumulate in non-recombining regions, e.g., in Y and W chromosomes [15–17], potentially including mutations that result in non-optimal expression [16]. In contrast, X and Z chromosomes can recombine in the homogametic sex and are thus less prone to degeneration. Sex-biased expression could thus result from degeneration, without involving sexually antagonistic selection, although this hypothesis has received little consideration [18,19]. For example, genes with differential expression on neo-sex chromosomes of the passerine bird *Sylvia communis* [20], and in the mating-type chromosomes in the hermaphroditic fungus *Neurospora tetrasperma* [21] were found to have elevated sequence divergence between alleles. While those studies pointed to sexual antagonism as the likely cause of differential expression between sexes and the relationship to sequence divergence between alleles, accumulated mutations with degenerative effects remain an alternative possibility.

Several different types of mutations can cause sequence degeneration and alter gene expression. Base pair substitutions and indels (insertion or deletion mutations) can change amino acid sequences which can affect gene expression through modulation of the mRNA translation [22], or disrupt promoter regions that impact transcriptional regulation [23]. Induction of early stop codons that truncate protein length can lead to post-transcriptional regulatory negative feedbacks upon expression (e.g. nonsense mediated decay; [24]). Transposable element insertion in upstream promoter regions, or internal to genes, has long been recognized to have effects on expression [25–29]. Epigenetic modifications, particularly cytosine methylation, contribute both to heterochromatin formation and elevating mutation rates that reduce GC content [30–32]; thus reduced GC content could represent a signature of methylation-induced gene silencing. Shorter introns are more efficient for correcting transcription [33], such that changes in introns can influence transcription rates, nuclear export, and transcript stability [34]. These forms of degenerative changes are expected to accumulate under the reduced selection efficacy in non-recombining regions. Yet, very few studies have addressed the relationship between sequence degeneration and differential gene expression on sex chromosomes [5,35,36].

The rarity of studies relating sequence degeneration to levels of differential gene expression is likely due to several major challenges they face. Studying degenerative processes requires assessing allele specific expression and comparing expression at various ages of sex chromosome divergence, including young sex chromosomes where recombination suppression events are recent. In commonly studied diploid organisms, however, sex chromosomes are old and already highly degenerated, where sex-linked gene expression can be impacted by gene presence/absence asymmetry between sex chromosomes and the resulting dosage compensation [37,38]. On the other hand, the investigation of the early stages of sex chromosome differentiation is rendered challenging by the lack of nucleotide differences which makes it difficult to confidently assigning transcripts to alleles in diploid organisms [39]. In addition, sex chromosomes are challenging to assemble in plants and animals, and the coverage-based method commonly used to identify sex-linked genes [19,40] is not applicable in young sex chromosome systems without much gene loss. For diploid systems with young sex chromosomes, sex-specific linkage maps are thus often needed to assign alleles to X/Z or Y/W chromosomes.

Fungi can provide valuable insights into the relationship between sequence degeneration and differential gene expression in sex or mating-type chromosomes, due to easy access to the haploid phase where alternate mating types are expressed, the existence of young events of recombination suppression in successive evolutionary strata, and the low potential for sexually antagonistic traits [41–44]. The anther-smut fungi, in the genus *Microbotryum*, undergo mating in the haploid phase via isogamous yeast-like cells of opposite mating types (a_1_ and a_2_), which can be cultured separately to analyze expression levels of alleles [45]. The species *Microbotryum lychnidis-dioicae,* causing anther-smut disease on the plant *Silene latifolia*, carry dimorphic mating-type chromosomes that have been assembled at the chromosome-level scale [42,43,46,47]. These mating-type chromosomes (a_1_, ~3.3Mb, and a_2_, ~4.0Mb, respectively) lack recombination across 90% of their length [46,47]. Importantly, evolutionary strata of different ages have been identified, i.e., regions with different levels of differentiation between mating types as a result of an expanding process of recombination suppression over the past ca. 1.5 million years [43,44]. The non-recombining regions of the mating-type chromosomes in *M. lychnidis-dioicae* are flanked by small recombining pseudo-autosomal regions (PARs).

In *M. lychnidis-dioicae*, the two possible causes leading to differential gene expression between the alternative haploid mating types are not equally probable, i.e. ‘mating-type antagonistic selection’ (*sensu* [48]) and differential sequence degeneration between alleles. The existence of genes under mating-type antagonistic selection in fungi would require fitness differences associated with mating-type dimorphic traits. However, previous studies on *M. lychnidis-dioicae* have shown that differences between the mating types, either developmental or ecological, are lacking outside of the immediate process of gamete fusion [41,49–52]. Moreover, a recent study on gene expression and positive selection detected no evidence for mating-type antagonistic selection [53]. This model system is therefore ideal to investigate the impact of degeneration on differential gene expression between chromosomes determining reproductive compatibility, notably without the confounding effect of sexual antagonism.

In this study, we therefore investigated whether mating-type specific differential gene expression was related to differences between alleles for various signatures of degeneration in the genome of *M. lychnidis-dioicae*. We then assessed the hypothesis that, for genes with differential expression between mating types, the alleles showing lower expression levels would have higher levels of degeneration footprints that are thought to be associated with disruption of gene expression. The investigated degeneration signatures included differences between alleles in the levels of non-synonymous sequence divergence, transposable element (TE) insertions, alteration of predicted protein length, intron content, and GC content (as a predicted consequence of epigenetic gene silencing) [27,28,33,54,55]. As prior work indicated that non-recombining regions of the mating-type chromosomes are enriched for signatures of sequence degeneration compared to autosomes [42], we also investigated whether differential gene expression varied among genomic compartments defined as autosomes, PARs, young evolutionary strata of the mating-type chromosomes (including previously identified red and green strata, [43]), and old evolutionary strata (blue, purple, orange and black strata, [43]).

## Results

### Allele identification and differential gene expression between a_1_ and a_2_ haploid genomes

Alleles of single-copy genes in *M. lychnidis-dioicae* were identified using the criterion of 1:1 reciprocal best BLASTp between a_1_ and a_2_ haploid genomes, based on the previously published genome assembly and gene annotation [43,44]. Protein sequence identity of >70% was used following evaluation of various identity thresholds (see details in Methods section). After filtering out TE-related gene sequences, we identified 371 single-copy allelic pairs in mating-type chromosomes and 9,025 in autosomes (S1 Table).

Using whole-genome RNA-seq data from separate a_1_ and a_2_ haploid mating-type cultures under low nutrient conditions that resemble the natural haploid growth environment [45,56], filtering for genes with significantly detectable expression recovered 8,549 single-copy genes for further analysis (342 on mating-type chromosomes and 8,207 on autosomes). The differential gene expression profile (i.e. (|Log2(a_1_/a_2_|) significantly greater than zero with false discovery rate (FDR) < 0.050, S1 Fig; S2 Table) revealed 392 genes (4.59% out of the 8,549 genes analyzed for expression) that were significantly more highly expressed in the a_1_ haploid culture, and 203 (2.37%) that were significantly more highly expressed in a_2_ haploid culture.

### Differential gene expression and multiple signatures of sequence degeneration

Regression analysis (generalized linear model, GLM) revealed that the degree of differential expression (DE) between allele pairs of the two haploid mating types significantly increased with increasing differences between alleles (using absolute values) in the various degeneration traits examined (Table 1). The significant main-effect predictors of differential expression included genomic compartment and differences between alleles in non-synonymous divergence (*dN*), transposable element (TE) insertion number within 20kb (up and downstream), intron content (proportional to coding sequence length), and overall GC content (GC0). Differences between alleles in predicted protein length was not a significant main-effect predictor but was strongly significant as an interaction term with genomic compartment and all other traits except intron content (Table 1). Differential expression indeed increased with differences between alleles in predicted protein length, but only in old evolutionary strata and when associated with higher differences between alleles in *dN*, TE content, and GC0 (Table 1; S2 Fig). Genes with differential expression between mating types showed significant enrichment in the old evolutionary strata compared to autosomes, but there was no enrichment in the young evolutionary strata or the PARs (Table 2). Similar patterns were observed for the comparisons in each of the a_1_ or a_2_ haploid genomes separately (S3 Table). Further *post hoc* assessments of degenerative traits are presented in the following sections, including whether difference between alleles is oriented such that the more affected allele is less expressed.

**Table 1.** Output of a reduced best-fit generalized linear model (GLM) with differential gene expression (|Log2(a_1_/a_2_)|) as the response variable and as predictable variables genomic compartment and various degeneration traits, i.e., non-synonymous mutation rate (*dN*), transposable element (TE) insertions, protein length, intron content and GC content. P values <0.05 are in bold. NA: not applicable.

| Predictable variables and interaction terms | GLM model output parameter |  |  |  |
| --- | --- | --- | --- | --- |
|  | Wald Chi-Square | Degree of freedom (df) | P value | Regression coefficient |
| (Intercept) | 496.78 | 1 | **<0.001** | NA |
| Compartment | 20.151 | 3 | **<0.001** | NA |
| $dN$ | 13.21 | 1 | **<0.001** | 5.081 |
| TE insertions | 8.405 | 1 | **0.004** | 0.044 |
| Protein length | 0.41 | 1 | 0.522 | 10.612 |
| Intron content | 10.209 | 1 | **0.001** | 0.768 |
| GC content | 4.233 | 1 | **0.040** | 0.499 |
| Compartment * Protein length | 24.662 | 3 | **<0.001** | NA |
| $dN$ * Protein length | 13.36 | 1 | **<0.001** | -50.726 |
| TE insertions * Protein Length | 8.398 | 1 | **0.004** | -0.37 |
| GC content * Protein length | 10.801 | 1 | **0.001** | -3.962 |

**Table 2.**
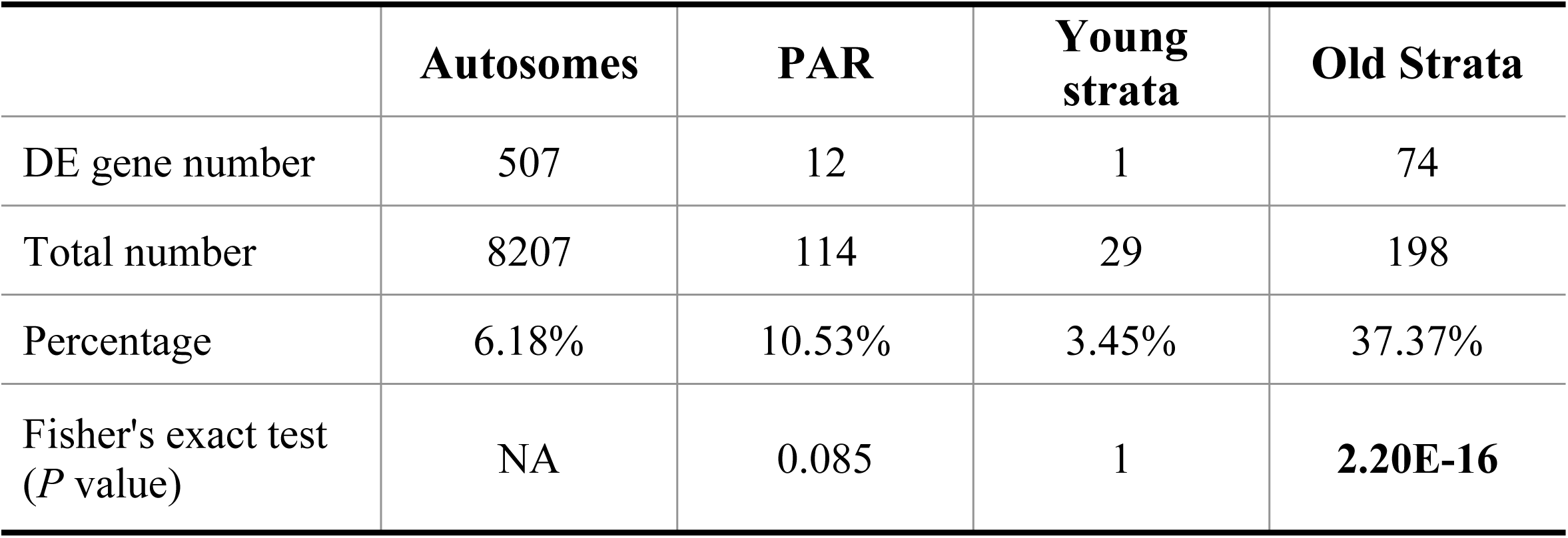
Numbers and percentages of genes with differential expression (DE) on the different genomic compartments on mating-type chromosomes and autosomes, and Fisher’s exact test for even distribution between DE genes on autosomes versus other genomic compartments, including pseudo-autosomal regions (PARs), young evolutionary strata (previously identified red and green strata; [43]) and old evolutionary strata (blue, purple, orange and black strata; [43]). P values <0.05 are in bold. NA: not applicable.

|  | **Autosomes** | **PAR** | **Young strata** | **Old Strata** |
| --- | --- | --- | --- | --- |
| DE gene number | 507 | 12 | 1 | 74 |
| Total number | 8207 | 114 | 29 | 198 |
| Percentage | 6.18% | 10.53% | 3.45% | 37.37% |
| Fisher's exact test (*P* value) | NA | 0.085 | 1 | **2.20E-16** |

### Relationship between differential expression and elevated substitution rates

Differentially expressed genes had greater sequence divergence between alleles than non-differentially expressed (non-DE) genes within genomic compartments, specifically within the old evolutionary strata of the mating-type chromosomes. DE genes had significantly higher non-synonymous mutation rate (*dN*) and synonymous mutation rate (*dS*) between alleles than non-DE genes within old evolutionary strata (Wilcoxon rank sum test for independent samples, *dN*: W = 1433, *P* < 0.001, *dS*: W = 1422, *P* < 0.001) (Fig 1A, S3 Fig, S4 Table). There was almost no sequence divergence (*dN* or *dS*) between alleles on either autosomes or PARs for DE or non-DE genes. The young evolutionary strata had only one DE gene, precluding comparison to non-DE genes within this compartment. The old strata pattern held for genes in a_1_ or a_2_ cells considered separately (S4A and S4B Fig). A tendency of higher *dN*/*dS* for DE genes compared to non-DE genes was not significant within old strata (W = 1946, *P* = 0.611) (S5 Fig).

**Fig 1.**
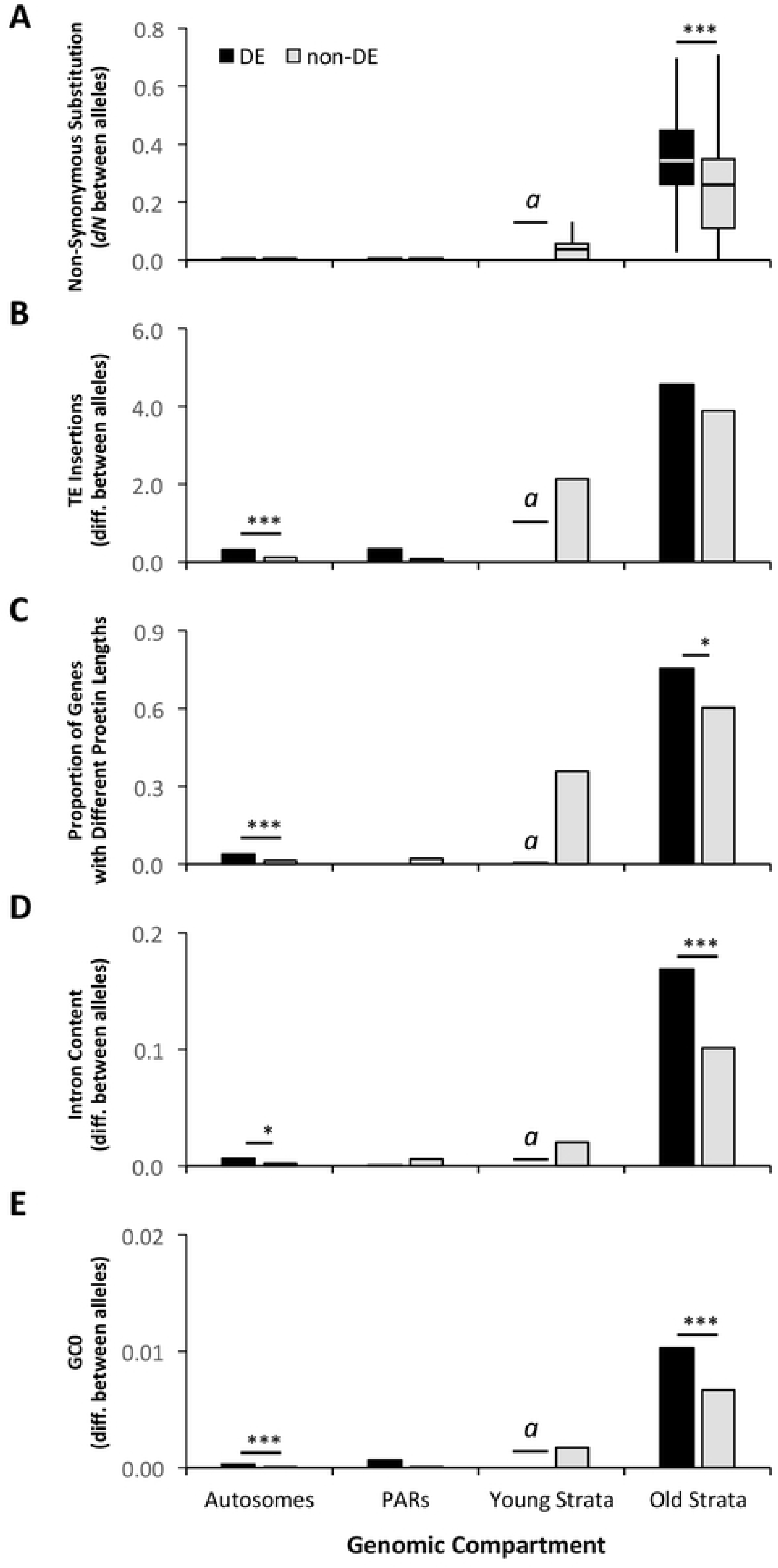
Comparisons of differentially expressed (DE) versus non-differentially expressed (non-DE) genes between mating types of *Microbotryum lychnidis-dioicae* for various degeneration-associated traits within genomic compartments. (A) Non-synonymous sequence divergence, *dN*, between alleles of DE and non-DE genes. (B) Transposable element (TE) insertion number differences between alleles within 20kb (up and downstream) of DE and non-DE genes. (C) Proportions of differentially expressed (DE) and non-differentially expressed (non-DE) genes with different protein lengths between alleles. (D) Intron content proportional differences between alleles of DE and non-DE genes. (E) Total GC content (GC0) proportional differences between alleles of DE and non-DE genes. Analyzed allele differences represent absolute values comparisons (i.e. unoriented with regard to allele expression levels). Comparisons in panels A, C-E reflect Wilcoxon rank sum tests; panel B reflects a two-proportion z-test. Significance levels shown as, ***: *P* < 0.001, *: *P* < 0.05; non-significant test results shown in Supplementary Tables S4, S6-S9. Genomic compartments include autosomes, pseudo-autosomal regions (PARs), young evolutionary strata (previously identified red and green strata; [43]) and old evolutionary strata (blue, purple, orange and black strata; [43]). The notation “*a*” indicates that the young evolutionary strata contained only one DE gene, precluding comparisons to non-DE genes within this compartment.

To test the hypothesis that the allele with lower expression would show a larger accumulation of non-synonymous changes than the allele with higher expression, each allele in *M. lychnidis-dioicae* was compared for sequence divergence with their ortholog in *M. lagerheimii*, which has retained largely collinear and homozygous mating-type chromosomes, as the inferred ancestral state in the *Microbotryum* genus [43,44]. Alleles in a_1_ haploid genome or in a_2_ haploid genome of *M. lychnidis-dioicae* were compared for *dN* divergence accordingly with alleles from the *M. lagerheimii* genome of the same mating type. Alleles having lower expression levels in *M. lychnidis-dioicae* did not have significantly greater *dN* divergence from their ortholog in *M. lagerheimii* than alleles with higher expression levels (W = 1,267, smallest *P* = 0.909) (S6 Fig, S5 Table).

### Relationship between differential expression and TE insertions

Differentially expressed genes were associated with greater differences between alleles for TE insertions (within 20kb up and downstream) than alleles of non-DE genes across genomic compartments. However, the difference was significant only in the autosomes (W = 313879, *P* < 0.001, Fig 1B), not in the PARs (W = 546, *P* < 0.192) or the old evolutionary strata (W = 4062, *P* = 0.173); the comparison was not possible in young evolutionary strata.

To test the hypothesis that the allele with less expression would show more TE insertions than the allele with higher expression, differences in TE insertions between alleles were calculated as the TE number for the allele with lower expression minus the TE number for the allele with higher expression; a positive value thus represented an excess of TEs in the less expressed allele. This oriented TE number difference between alleles was tested as a predictor of the expression ratio |Log2(a_1_/a_2_)| using a sliding window approach with a 15kb window size overlapping by 5kb. Among DE genes, oriented TE insertion difference was a significant predictor of the express ratio only in the window covering from 10kb upstream to the gene (Fig 2A, S7 Fig); alleles with more TE insertions having reduced expression (for this window, Wald X^2^ = 6.674, *P* = 0.010, statistics of remaining windows in S6 Table). Among non-DE genes, none of the windows was a significant predictor of variation in the expression ratio (S6 Table).

**Fig 2.**
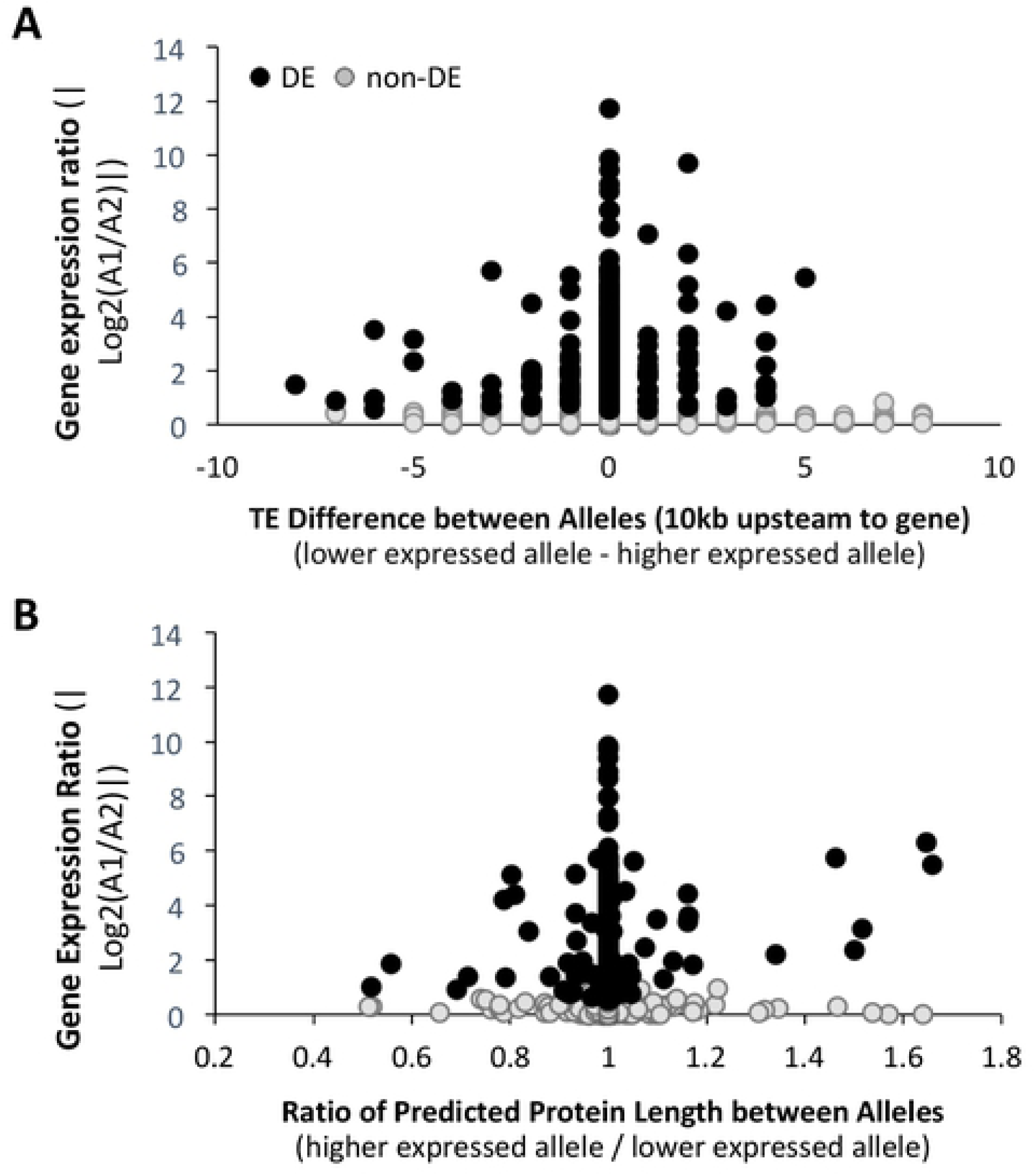
Significant predictors of the degree of differential expression between mating types of *Microbotryum lychnidis-dioicae* testing directional effects of degeneration-associated traits. (**A**) Relationship of expression ratio to oriented TE insertion differences in the region from 10kb upstream to the gene, where the trait was calculated as TE number for the allele with lower expression minus the TE number for the higher expressed allele; a positive value thus represented an excess of TEs in the lower expressed allele. (**B**) Relationship of expression ratio to oriented predicted protein length differences, where the trait was calculated as the ratio for the allele with higher expression divided by the allele with lower expression; a larger ratio thus represented a shorter length for the allele with lower expression.

### Relationship between differential expression and differences in predicted protein length

Differential gene expression was associated with the mutational changes that affect the predicted protein length, including altered stop codon positions, indels, and indels causing frameshifts. Within genomic compartment, alternate alleles of DE genes were significantly more likely to produce proteins of different lengths than alleles of non-DE genes, particularly within the old evolutionary strata (Two proportion Z test, z = 2.186, *P* = 0.029) and autosomes (z = 4.64, *P* = 8.78e-06, Fig 1C, S7 Table); there were too few DE genes on PARs and young evolutionary strata for statistical comparisons.

The various types of mutational changes that caused protein length variation between alleles differed between DE and non-DE genes, as well as among genomic compartments. Among the 258 genes with different protein sequence lengths between alleles, all had indels. However, DE genes in the old evolutionary strata and autosomes had significantly more indels than non-DE genes; old strata mean indel number differed between alleles by 2.64 for DE genes and by 1.85 for non-DE genes (W = 2453.5, *P* = 0.013), and in autosomes alleles differed by a mean of 1.19 indels for DE genes and 1.03 for non-DE genes (W = 490.5, *P* = 0.025, S8A Fig); PARs and young evolutionary strata could not be analyzed.

Similarly, differences in the positions of stop codons contributed to protein length variation more for DE genes than non-DE genes. Among genes with different protein lengths between alleles in the old evolutionary strata, 44.6% (N = 56) of DE genes had different stop codon positions between alleles, which was significantly higher than the 24.0% (N = 75) of non-DE genes (two-proportion z-test, *P* = 0.018, S7B Fig). Similarly, DE genes in the autosomes were marginally significantly more likely to have different stop codon positions between alleles than non-DE genes, with 33.3% (N = 21) vs 10% (N = 40), respectively (*P* = 0.057, S8B Fig). Only three frameshift mutations were observed among the 258 of genes examined with different protein/coding sequence lengths, and thus frameshifts were not distinguishing features of DE versus non-DE genes.

To test the hypothesis that the allele with less expression would show a truncation of protein length compared to the allele with greater expression (i.e. by early stop codons or deletions), differences in protein length between alleles were calculated as the ratio for the allele with higher expression divided by the allele with lower expression; a larger ratio thus represented a shorter length for the allele with lower expression. Among DE genes, this oriented metric of protein length differences was a significant predictor of the differential expression degree as the ratio |Log2(a_1_/a_2_)|, with alleles producing shorter proteins being less expressed (Wald X^2^ = 19.326, two-tailed *P* < 0.001, Fig 2B). No significant relationship to expression level ratio was found for the length ratios of non-DE genes (Wald X^2^ = 0.222, *P* = 0.638, Fig 2B).

### Relationship between differential expression and intron content

Differential gene expression was associated with differences between alleles in intron content, considering lower intron content to be favored by selection [33]. There were significantly greater intron content differences between alleles for DE genes than for non-DE genes; considering the ratio of intron to coding sequence lengths, alleles of DE gene overall differed on average by 0.008 and alleles of non-DE genes differed by 0.002 (W = 2102758, *P* < 0.001). Alleles differed in intron content more for DE than non-DE genes within the autosomes (W = 1920124, *P* = 0.033) and old evolutionary strata (W = 3205, *P* = 0.001) (Fig 1D, S8 Table), but not within the PARs (W = 605, *P* = 0.888); the comparison in young evolutionary strata was not possible.

To test the hypothesis that the allele with lower expression would show a greater intron content as a signature of degeneration, differences between alleles were calculated as the value for the less expressed allele minus the value for the more expressed allele; a positive value thus represented greater intron content for the less expressed allele. This oriented metric of intron content differences between alleles was not a significant predictor of differential expression level among DE genes (Wald X^2^ = 0.350, *P* = 0.554), or among non-DE genes (Wald X^2^ = 0.216, *P* = 0.642).

### Relationship between differential expression and GC content

Consistent with gene silencing by cytosine methylation contributing to decreased GC content [30,32,57], DE genes had significantly greater overall GC0 differences between their alleles than non-DE genes within the autosomes (W = 1907831, *P* < 0.001) and old evolutionary strata (W = 3010, *P* < 0.001) (Fig 1E). The comparison within the PARs was not significant (W = 578, *P* = 0.318); the comparison for young evolutionary strata was not possible. Analysis of third codon position GC3 provided similar patterns and levels of significance (S9 Fig, S9 Table).

To test the hypothesis that the allele with lower expression would show lower GC content than the allele with higher expression, GC0 or GC3 differences between alleles were calculated as the value for allele with higher expression minus the value for allele with lower expression; a positive value thus represented reduced GC content for the allele with lower expression. Among DE genes, neither the oriented GC0 or GC3 differences between alleles were significant predictors of the level of differential expression (GC0: Wald X^2^ = 1.039, *P* = 0.308, and GC3: Wald X^2^ = 2.226, *P* = 0.136).

## Discussion

Genomic regions controlling mating compatibility, whether non-recombining sex or mating-type chromosomes, have been subject of intense research. This is because these regions determine traits essential for fitness and because of their rapid evolutionary dynamics. Sequence degeneration of non-recombining regions has been linked to major genomic features such as chromosomal heteromorphism, dosage compensation or gene trafficking between non-recombining regions and autosomes [14,37]. To our knowledge, our study is the first to reveal an association between a variety of degeneration measures and differential gene expression, doing so in the anther-smut fungus *M. lychnidis-dioicae* where antagonistic selection is unlikely [41,43,44,52,53], and thus does not constitute a confounding factor. Genes differentially expressed between the haploid mating types were enriched on the oldest strata of the mating-type chromosomes and displayed various forms of sequence (*dN*, *dS*, or GC content) or structural (TE insertions, introns content, or protein length) heterozygosity at levels higher than non-differentially expressed genes. These results across various forms of mutational change and genomic compartments show importantly that differential gene expression is strongly associated with sequence degeneration, and differential expression should therefore not be systematically interpreted as a sign of antagonistic selection.

### Differential gene expression between haploid mating types

The proportion of genes with differential expression between haploid mating types of *M. lychnidis-dioicae* was low, which is consistent with expectations based upon the lack of dimorphism between mating types. The number of DE genes was slightly higher than in a previous study based on the same dataset [42], likely due to an improved genome assembly and non-recombining region identification. The 2.4~4.6% of genes with differential expression between mating types of *M. lychnidis-dioicae* was similar to plant and animal non-reproductive tissues, e.g. liver, spleen, leaves, roots [9,58–64]. However, this is much lower than in reproductive tissues (e.g. ovaries or testes) of most animals and plants [5,6]. There seems to be an overall positive relationship between proportions of sex-biased genes and levels of sexual dimorphism across taxa [65], suggesting a possible correlation between phenotypic difference between sexes and underlying transcriptional architecture. The proportion of sex-biased genes was lower than the 12% found in the haploid sexes in the brown algae *Ectocarpus* which has a low level of sexual dimorphism [66]. Among fungi, 3~4% of genes had differential expression between mating types in the fungus *Neurospora tetrasperma* [21]. Therefore, the low percentage of genes having differential expression between mating types in *M. lychnidis-dioicae* is consistent with its life history having no ‘female’ or ‘male’ functions (isogamous gametes) and mating type being controlled at the haploid stage without morphological or ecological differences that might provide different trait optima between mating types [41,53,67].

Differentially expressed (DE) genes were enriched in the mating-type chromosomes of *M. lychnidis-dioicae*, which is also consistent with studies in animals and plants having differentiated sex chromosomes (reviewed by [5]). Similarly, in the fungus *N. tetrasperma*, DE genes were more frequently detected on mating-type chromosomes [21]. In animals and plants, sexually antagonistic selection is accepted as an important force that can explain the stepwise recombination suppression in sex chromosomes and formation of evolutionary strata. Indeed, linkage of sexually-antagonistic genes to reproductive compatibility loci is considered fundamental to the resolution of sexual conflict by allowing for sex-specific or sex-biased gene expression [2,68–70]. Sex-biased gene expression is often considered the product of sexually antagonistic selection across a broad range of taxa [5,6,20,21,71]. However, decades of research have uncovered little genetic evidence directly supporting sexually antagonistic selection as being the driving force for evolutionary strata of recombination suppression or for sex-biased expression outside of the reproductive tissues themselves [7,19,58,72].

Our results, however, suggest a broader view of evolutionary forces, aside from sexual antagonism, that can explain the occurrence of DE genes and their enrichment on chromosomes determining reproductive compatibility. The role of sequence degeneration in differential expression has so far been largely understudied but is likely the primary and powerful factor in *M. lychnidis-dioicae*. The lack of female and male functions in *Microbotryum* fungi and a haploid phase with very limited differences between mating types [41], likely explains the absence of genes experiencing mating-type antagonistic selection [53]. In contrast, gene degeneration on non-recombining sex or mating-type chromosomes occurs commonly among eukaryotic taxa due to the combined influences of reduction in effective population size (*N_e_*), Hill-Robertson interference and sheltering effects in diploid organisms [2,5,42,68–70]. Sequence degeneration is therefore a generally expected phenomenon in non-recombining regions, and the resulting mutation accumulation may generate contrasting expression levels between differently affected alleles. Certainly, sexually antagonistic selection and degeneration are not mutually exclusives processes, and a general relationship between the amounts of sexual dimorphism and of DE genes [5,6] supports the role for sexual antagonisms. However, several types of degeneration are found in *M. lychnidis-dioicae*, in the absence of sexual antagonism, suggesting the potential for similar in patterns associated with DE genes across diverse types of organisms.

### Various forms of degeneration

The properties of non-recombining regions that reduce the efficiency of selection (reduced *N_e_*, hitchhiking and sheltering) can lead to the fixation of various mutations having degenerative effects, several of which were significant predictors in the overall regression model of differential expression between mating types of *M. lychnidis-dioicae*. Some signatures of degeneration can be directly connected to mechanisms known to reduce transcription levels. In particular, transposable element (TE) insertion into genes or upstream have long been recognized to alter gene expression [29,73]. TEs can disrupt promoter regions or other regulatory sequences internal to genes [27,28]. In addition, epigenetic silencing, as a defense against TE proliferation, can tighten local chromatin structure and inhibit access of transcriptional machinery [74,75]. In this study, the differences between alleles in DE genes in the number of TE insertions were a significant predictor of differential expression. Consistent with a direct effect upon differential expression, the relative excess of TE insertions between alleles, specifically upstream of genes, was associated with a lower expression level between alleles of DE genes. Similarly, the introduction of early stop or non-sense codons is expected to reduce expression. Transcripts from alleles with premature stop codons are affected by nonsense mediated decay, involving degradation of mRNA and further components of the RNAi pathway than down-regulate expression [54]. We found that alternative alleles were more likely to have differing stop codon positions in DE genes than in non-DE genes, and, importantly, that the differential expression was explained by the shorter allele having lower expression. This suggests that gene truncation, by the gain of premature stop codons rather than the possibility of stop codon losses and read-through transcripts, leads to differential expression. Both TEs and premature stop codons appear to be important mutations changes that affect differential expression between alleles.

Other characteristics of DE genes in *M. lychnidis-dioicae* are less mechanistically tied to changes in expression levels but are nevertheless associated with sequence degeneration, directly or indirectly. Most important among these characteristics was the degree of sequence divergence between alleles. Alleles of DE genes were distinguished by markedly more non-synonymous and synonymous base pair differences than alleles of non-DE genes, sequence divergence being a positive predictor of the degree of differential gene expression. Similar results were demonstrated in the anisogamous, hermaphroditic ascomycete *N. tetrasperma*, showing that differential gene expression was positively correlated with sequence divergence between alleles of genes on mating-type chromosomes [21]. Gene profiles in *N. tetrasperma* differed in their mating-type-specific expression depending upon whether female or male reproduction was being induced by the culture conditions, drawing analogy to sexual dimorphism commonly found in animals and plants [21]. However, *Microbotryum* fungi do not have such male or female functions or ecological differences between mating types, and therefore such an adaptive explanation for DE genes is improbable.

The remaining signatures of degeneration were most informative of how genes might evolve as a consequence of differential expression. In general, gene expression levels are expected to positively correlate with the strength of selection [55]; stronger selection and higher expression levels have been correlated with reduced intron content [33]. It is also known for many eukaryotes that introns can affect gene expression without functioning as a binding site for transcription factors [76], by for example influencing the rate of transcription, nuclear export and transcript stability [34]. Our data show that differences between alleles in intron content positively predict differential expression level, which is consistent with these intron effects upon gene expression, although the directional hypothesis testing was not significant. Additionally, our data showed changes in GC content that are consistent with the consequences of suppression of gene expression, specifically as predicted to result from epigenetic silencing through cytosine methylation. Two important drivers of GC content changes are biased gene conversion and methylcytosine-driven C-to-T mutation rates [31,32]; however, gene conversion that relies on meiotic pairing is unlikely to cause the different mutation rates between alleles of single copy genes [77]. While differences between alleles in GC content was a modestly significant predictors of differential expression level compared to some other traits in the overall regression model, we also found alleles of DE genes differed more in GC content than non-DE genes in the old evolutionary strata. Methylation-mediated gene silencing has been presented as one of the degeneration types following recombination suppression on sex chromosome of *Drosophila albomicans* [16,78]. Those studies indicate that reduced chromosome-wide gene expression occurred as a first step in degeneration between non-recombining regions of a young sex chromosome system [16,78]. While our findings are in support of the hypothesis for methylation-mediated effects on differential expression, methylome analyses across the *M. lychnidis-dioicae* genome should be investigated in more details.

### Degeneration across genomic compartments

The different forms of genetic degeneration in *M. lychnidis-dioicae* were not equally represented among genomic compartments, perhaps reflecting the history of recombination suppression. In this system, enrichment of DE genes on the mating-type chromosomes is unlikely to be due to antagonistic selection, but rather to this region preserving heterozygosity in general. As a matter of fact, enrichment of DE genes was significant only in the old evolutionary strata and not in the younger strata, indicating it is a consequence and not a driver of recombination suppression.

Mating between different haploid sexes or mating types ensures that all diploids are heterogametic [79], and it has long been recognized that regions linked to mating type can preserve heterozygosity [80]. In *M. lychnidis-dioicae*, the large non-recombining regions are in fact highly heterozygous [43]. In contrast, the autosomes and PARs are largely homozygous, due to the selfing mating system of *M. lychnidis-dioicae* [41,47]. Consistent with mating-type linkage preserving heterozygosity, nearly the full range of mutational changes or footprints of degeneration showed lowest levels in the autosomes and PARs and increasing through the young evolutionary strata to highest levels in the old evolutionary strata. Traits following this pattern included differences between alleles in TE accumulation, base pair substitutions, protein length, intron content and GC percentage. In most cases, these effects increased also for non-DE genes in the non-recombining region of the mating-type chromosomes but to lower degrees than DE genes. Importantly, however, comparisons within genomic compartments repeatedly showed that allele distinguishing mutations occurred more in association with DE genes than non-DE genes or in the manner positively associated with levels of differential expression. Therefore, strong evidence is shown for these degenerative changes being directly related to changes in expression levels between alleles.

These degeneration patterns are consistent with prior studies on *M. lychnidis-dioicae* showing the existence of deleterious recessive alleles that are linked to mating type and preventing haploid growth [81,82], which may reflect outcomes of the mutational accumulation described here. The recent discovery of multiple independent mating-type linkage events across the *Microbotryum* genus [44] should allow further assessment of mutation accumulation and its consequences for gene functions. Finally, it should be noted that the acquisition of one type of degenerative mutation, particularly where expression is reduced, may relax selection and favor the accumulation of additional sequence changes in that allele. Whether such feedback or cascade dynamics further drive allele denervation should be addressed in further studies.

### Insights across sexual eukaryotes

*Microbotryum*, and fungi in general, provide powerful models to investigate the genomics of sequence degeneration and differential gene expression between sex-related chromosomes. The ability to culture haploid genotypes helps to overcome issues of allele-specific expression encountered in diploid systems. The genomes are small and readily assembled at the near-chromosome level. Also, some pathogenic fungi like *Microbotryum* exhibit an obligately sexual life cycle, where meiosis and haploid mating are required upon each instance of disease transmission [41]. Furthermore, fungi often have a restricted range of evolutionary forces related to a sexual life cycle (i.e. without sexually antagonistic selection and without asymmetrical sheltering resulting from the haploid mating; [79]). Therefore, finding traits in common between *Microbotryum* and familiar plant and animal systems help to identify fundamental evolutionarily mechanisms for the genomics of reproductive compatibility [83]. In particular, the recent demonstration of multiple evolutionary strata on the mating-type chromosomes of *M. lychnidis-dioicae* and other *Microbotryum* species [43,44] reinforces the idea that mechanisms aside from sexual antagonism are sufficient to drive such patterns of recombination cessation [84]. The evolutionary strata in *Microbotryum* provide remarkable opportunities to study the accumulation of sequence divergence traits over time. The current study further reveals differential expression of genes as a common feature of non-recombining mating-type chromosomes and sex chromosomes, where, in the absence of sexual antagonism, a major role for degenerative mutations is indicated.

### Conclusions

Our findings on differential gene expression being associated with various types of sequence degeneration, and likely being a consequence of those, shed new lights on how differential gene expression can evolve. In animals and plants, it is widely accepted that differential gene expression on sex chromosomes is associated with sexually antagonistic selection [37,85]. Our study shows that, in systems where sexual antagonistic selection is unlikely to occur, sequence degeneration might readily lead to differential gene expression. Furthermore, the genes with differential expression were highly enriched on mating-type chromosomes, similar to diverse organisms where in contrast the separate sex functions have been cited as the cause. We further found strong and consistent evidence of differential gene expression and its association with various types of mutational changes, in particular TE insertions and premature stop codons, as well as high levels of base pair substitutions, indels, intron and GC content that distinguish alleles of DE genes. We show that sequence degeneration in fact largely characterizes DE genes identified between fungal mating types, in the absence of sexually antagonistic selection, and only on the old evolutionary strata. Our results help to uncover important patterns of gene evolution relevant to a broad range of taxa where reproductive compatibility is determined in extensive regions of recombination suppression.

## Materials and Methods

### Allele identification between a_1_ and a_2_ haploid genomes

In order to quantify differentially expressed genes between the two haploid genomes, the alleles between a_1_ and a_2_ haploid genomes need to be identified for those genes. The genome assembly and annotation of the same strain of *M. lychnidis-dioicae* have been published [43]. To identify 1:1 single copy homologs in each haploid genome, the Reciprocal Best BLAST(p) Hits (RBBH) python script (github.com/peterjc/galaxy_blast/tree/master/tools/blast_rbh) was applied [86], with 50 percentage of length coverage. RBBH scripts also identified paralogs within each haploid genome. A number of protein sequence alignment identity thresholds were tested, in order to identify for the best strategy of maximizing the number of allele pair identification on the non-recombining regions and while avoiding spurious BLAST results with low identity percent. Increasing the percent of protein sequence identity threshold from >70% to >85% resulted in a decrease from 12.2% to 9.9% of single-copy genes on the mating-type chromosomes being identified as differentially expressed genes (detailed below), while decreasing the threshold from >70% to >30% resulted in only a marginal increase from 12.2% to 12.7%. The change in the percentages of identified alleles that were differentially expressed on autosomes was negligible, being 1.0%, 1.1% and 1.1% respectively for 80%, 70% and 30% thresholds (S10 Fig). Therefore, the threshold of >70% protein sequence identity was used. To avoid potential bias due to paralogs for identifying differential gene expression and other downstream analysis, genes with paralogs within each haploid genome were filtered out and only single-copy allele pairs were retained for downstream analysis. Genes were located to genomic compartments, including autosomes, pseudo-autosomal regions (PARs), young evolutionary strata of the mating type chromosomes (including previously identified red and green strata; [43]) and old evolutionary strata (blue, purple, orange and black strata; [43]).

### Transposable element filtering

Transposable element (TE) annotation of both haploid genomes of *M. lychnidis-dioicae* was published previously [87], and was used for analysis in this study. The coding sequence of each gene from both a_1_ and a_2_ haploid genomes was search by BLAST(n) against the published annotated TE consensus sequences of the same species, and alignment >80 percent of query coverage (coding sequences) was used for identifications of TEs. The BLASTn output was parsed using BASH scripts, and the coding sequences identified as TEs were removed from the gene list for all further downstream analysis.

### Identification of differentially expressed genes

RNAseq data and summary statistics of the datasets were described previously [45], and the raw data of haploid culture growing separately in water agar conditions were downloaded from the deposited NCBI database (https://trace.ncbi.nlm.nih.gov/Traces/study/?acc=+PRJNA246470&go=go). First, the RNAseq raw reads were quality assessed using FastQC v0.11.2 (https://www.bioinformatics.babraham.ac.uk/projects/fastqc/), and quality trimmed using Trimmomatic v0.33 with default parameters for paired-end reads [88]. We filtered reads containing adaptor sequences and trimmed reads if the sliding window average Phred score over four bases was < 15 or if the leading/trailing bases had a Phred score < 3. Reads were then removed post filtering if either read pair was < 36 bases.

To avoid possible bias for calling differential gene expression due to differences in homolog length between a_1_ and a_2_, gaps differing between alleles by greater than 3bp were trimmed to keep the same length, using published custom Python script [89]. This trimming includes the gaps from the ends of the alignment and inside the alignment, with inside gaps starting with the closest to the end of the alignment (greater than the minimum gap size) until there are no gaps larger than minimum gap size [89]. The trimmed allele pairs with equal length were used for read mapping and calling differential gene expression.

To quantify gene expression, we mapped the trimmed reads of haploid samples to the trimmed homolog sequences of each haploid genome respectively with Kallisto v.0.43.0 [90]. Read counts of the output from Kallisto mapping (e.g. using pseudo-alignment) were imported for gene expression analysis in EdgeR v3.4 [91,92]. We filtered low counts and kept genes with average Log(CPM) > 0 per sample, and CPM > 1 in half of the samples per haploid culture. We then normalized the expression by trimmed mean of M values (TMM). We explored the libraries of both haploid cultures in two dimensions using multi-dimensional scaling (MDS) plots (S11 Fig). Normalized expression counts for each sample were used to calculate differential expression between mating types using standard measures. We first identified genes with differential expression between mating types based on overall expression of the comparison group, and using Benjamini-Hochberg correction for multiple-testing with false discovery rate (FDR) of 5%. Differential expression between mating types was classified into four categories of fold changes, namely 2 (low), 2-4 (mild), 4-8 (high), and > 8 (very high), and expressed as log_2_ ratio of a_1_-to-a_2_ expression (which has negative values for genes with higher a_2_ expression and positive values for higher a_1_ expression). As suggested by [93], fold changes > 0 will be interpreted throughout, because we are working on haploid cell cultures and there are no possible scaling nor allometry issues due to whole-body sampling. Thus, unless stated otherwise, both conditions FDR < 0.05 and |log_2_FC| > 0 will be met when calling mating-type bias.

The classification of genes as having differential expression between mating types or the absolute values of gene expression ratio |Log2(a_1_/a_2_)| was used to assess relationships to various forms of mutational changes. Generalized linear model (GLM) analysis was used to assess the predictors of absolute values of expression ratio |Log2(a_1_/a_2_)|, with main effect variables and all two-way interactions terms for genomic compartments and the absolute value of differences between alleles for sequence divergence (*dN*), transposable element insertions number within 20kb (up and downstream), predicted protein length, intron content and GC content. The absolute value of the differences between alleles was calculated for each trait as detailed below. Model family comparison was based upon minimizing Akaike’s Information Criterion and over/under-dispersion using ratio of deviance/df; Tweedie, power 1.7 (approaching gamma distribution) provided the best available fit for the expression ratio response variable. A best fit model was selected using stepwise model selection, following removal of non-significant interaction terms. Other *post hoc* tests evaluating individual degeneration trait are described below. All statistical analyses were conducted in SPSS v23 [94] and R v3.4.3 [95].

### Relationship between differential expression and elevated substitution rates

Pairs of alleles between a_1_ and a_2_ mating types were aligned with PRANK (v170427) using the codon model [96]. Each pair of allele alignment was then analyzed with codeml in PAML [97] (runmode -2) to calculate the number of nonsynonymous substitutions per nonsynonymous site (*dN*), the number of synonymous substitutions per synonymous site (*dS*), and the ratio of the two (*dN*/*dS*), the latter excluding genes with *dS* value of zero. We then compared sequence divergence between alleles using non-parametric Wilcoxon rank sum tests for DE versus non-DE genes within genomic compartments.

Also, the allele sequences were compared between *M. lychnidis-dioicae* and their orthologs in *M. lagerheimii*, which has retained largely collinear and recombining mating-type chromosomes, as the inferred ancestral state in the *Microbotryum* genus [43,44]. The single-copy orthologs for a_1_ or a_2_ genomes between *M. lychnidis-dioicea* and *M. lagerheimii* were identified using RBBH with 70 percent protein sequence coverage identity (github.com/peterjc/galaxy_blast/tree/master/tools/blast_rbh, [86]). Wilcoxon rank sum test was used to assess *dN* between orthlogs in *M. lychnidis-dioicea* and *M. lagerheimii* to evaluate the hypothesis that the alleles with lower expression levels would have greater sequence divergence.

### Relationship between differential expression and TE insertions

The TE annotation of the *M. lychnidis-dioicae* genome published previously [87] was used for the analysis in this study. First, the TE insertion sites were assessed for each given focal gene, upstream 0-5k, 5-10kb, 10-15kb, 15-20kb distance intervals, and downstream 0-5kb, 5-10kb, 10-15kb and 15-20kb distance intervals using Bedtools window function for each indicated distance window (https://bedtools.readthedocs.io/en/latest/content/tools/window.html). Both annotation GFF3 files of gene models and TE annotations of *M. lychnidis-dioicae* were provided as input files. The output files were parsed using Bash scripts. Wilcoxon rank sum tests were used to compare TE insertions for DE and non-DE genes within genomic compartments.

Also, a limited GLM model was used to assess the hypothesized directional association of TE insertions and reducing allele expression (|Log2(a_1_/a_2_)|); this model contained genomic compartment and oriented TE differences between alleles as main effects and their interaction term. Oriented TE differences between alleles were calculated as the TE number for the allele with lower expression minus the TE number for the higher expressed allele; a positive value thus represented an excess of TEs in the lower expressed allele. A sliding window approach was used with a window size of three adjacent intervals, progressing from upstream to downstream of the genes.

### Relationship between differential expression and differences in predicted protein length

We first verified whether there was bias in the gene prediction model across genomic compartments, using the ratio of predicted coding sequence length divided by three times protein sequence length, assessed using linear regression model. Coding sequencing length divided by the length of predicted protein multiplied by 3 was consistently close to 1 and did not differ among genomic compartments (autosome, PAR, young strata and old evolutionary strata; Linear model, R^2^ = -5.50e-05, F-statistic = 0.869, P= 0.530, S12 Fig). We therefore calculated the ratio of predicted protein length between allele pairs, and compared the proportions of genes in DE and non-DE categories that had unequal lengths using two-proportion Z test for genes within genomic compartments. The mutational causes of unequal protein lengths was assessed by manually quantifying premature stop codons or indels using Geneious v8.1.7 [98]. A limited GLM model was used to assess the hypothesized directional association of protein truncation and reducing allele expression (|Log2(a_1_/a_2_)|); this model contained genomic compartment and oriented predicted protein length differences between alleles as main effects and their interaction term. Oriented predicted protein length differences between alleles were calculated as the ratio for the allele with higher expression divided by the allele with lower expression; a larger ratio thus represented a shorter length for the allele with lower expression.

### Relationship between differential expression and intron content

Using the published annotation gene models and coding sequences, we extracted the intron number and mean intron length information from the annotation gff3 file, using Perl script (https://bioops.info/2012/11/intron-size-gff3-perl/). We investigated the proportional differences of the intron content for both DE and non-DE genes within genomic compartments using Wilcoxon rank sum test. We also used a limited GLM model to test the hypothesized directional association of greater intron content and reducing allele expression (|Log2(a_1_/a_2_)|); this model contained genomic compartments and oriented intron content differences between alleles as main effects and their interaction term. Oriented intron content differences between alleles were calculated as the value for the lower expressed allele minus the value for the higher expressed allele; a positive value thus represented greater intron content for the lower expressed allele.

### Relationship between differential expression and GC content

We calculated the total GC percentage (GC0) and the GC percentage at the third position of aminol acid (GC3) for alleles of each gene coding sequence using homemade awk scripts. We investigated the differences of GC0 and GC3 for both DE and non-DE genes within genomic compartments using Wilcoxon rank sum test. We also used a limited GLM model to test the hypothesized directional association of reduced GC content and reducing allele expression (|Log2(a_1_/a_2_)|); this model contained genomic compartments and oriented GC content differences between alleles as main effects and their interaction term. Oriented GC content differences between alleles were calculated as the value for the allele with higher expression minus the value for the allele with lower expression; a positive value thus representing reduced GC content for the allele with lower expression.

### Ethics Statement

N/A

## Supporting Information Legends

**S1 Table. Number of single-copy genes with alleles in both a_1_ and a_2_ haploid genomes, with 70% protein sequence identity detection using reciprocal best BLASTp hits, before and after filtering out genes with transposable element (TE)-related functions.**

Filtering removed 192 paralogous genes within each haploid genome and genes with TE-related functions, including 1750 and 1819 from a_1_ and a_2_ haploid genome, respectively.

**S2 Table. Identification of differentially expressed (DE) genes with either a_1_-biased or a_2_-biased expression (i.e., higher expression in a_1_ or a_2_, respectively) under various Log2(a_1_/a_2_) criteria.**

**S3 Table. Numbers and percentages of genes with differential expression (DE) within genomic compartments, including autosomes pseudo-autosomal regions (PARs), young evolutionary strata (previously identified red and green strata;** [43]**) and old evolutionary strata (blue, purple, orange and black strata;** [43]).

Chi-squared test was used to assess whether DE genes were non-randomly distributed between autosomes versus the genomic compartments on mating type chromosomes (MAT). *P* values <0.05 are in bold. NA: not applicable. The young strata only had one DE gene, so statistic comparisons could not be performed for these strata.

**S4 Table. Wilcoxon rank sum test statistics for comparisons of mean non-synonymous mutation rate (*dN*, A) and synonymous mutation rate (*dS*, B) of differentially expressed genes (DE) versus non-differentially expressed genes (non-DE) within genomic compartments, including autosomes pseudo-autosomal regions (PARs), young evolutionary strata (previously identified red and green strata;** [43]**) and old evolutionary strata (blue, purple, orange and black strata;** [43]).

*P* values <0.05 are in bold. NA: not applicable. NA: not applicable, as the young strata only had one DE gene, statistical comparisons could not be performed for this compartment.

**S5 Table. Wilcoxon rank sum test statistics for comparisons of divergence of alleles in *Microbotryum lychnidis-dioicae* from orthologs in *M. lagerheimii*.**

This test assessed whether the allele with lower expression in *M. lychnidis-dioicae* was more divergent from orthologs in *M. lagerheimii* than the alleles with higher expression in *M. lychnidis-dioicae*, considering non-synonymous mutation rate (*dN*), synonymous mutation rate (*dS*), and the ratio (*dN/dS*) within genomic compartments, including autosomes pseudo-autosomal regions (PARs), young evolutionary strata (previously identified red and green strata; [43]) and old evolutionary strata (blue, purple, orange and black strata; [43]). We calculated these substitution rates for a_1_ and a_2_ alleles between these two species separately. NA: not applicable, as the young strata only had one DE gene, statistical comparisons could not be performed for this compartment.

**S6 Table. Wilcoxon rank sum test statistics for comparisons of unoriented transposable elements (TEs) insertion differences between alleles (within 20kb up and downstream) of differentially expressed (DE) genes versus non-differentially expressed (non-DE) genes within genomic compartments, including autosomes pseudo-autosomal regions (PARs), young evolutionary strata (previously identified red and green strata;** [43]**) and old evolutionary strata (blue, purple, orange and black strata;** [43]).

P-values <0.05 are in bold. NA: not applicable, as the young strata only had one DE gene, statistical comparisons could not be performed for this compartment.

**S7 Table. Two proportion Z test for comparisons of unoriented protein length difference between alleles of differentially expressed (DE) genes versus non-differentially expressed (non-DE) genes within genomic compartments, including autosomes pseudo-autosomal regions (PARs), young evolutionary strata (previously identified red and green strata;** [43]**) and old evolutionary strata (blue, purple, orange and black strata;** [43]).

*P* values <0.05 are in bold. NA: not applicable, as proportions * sample size was less than 5 for the PARs and young strata, statistical comparisons could not be performed for these compartments.

**S8 Table. Wilcoxon rank sum test statistics for comparisons of unoriented differences in intron content between alleles of differentially expressed (DE) versus non-differentially expressed (non-DE) genes within genomic compartments, including autosomes pseudo-autosomal regions (PARs), young evolutionary strata (previously identified red and green strata;** [43]**) and old evolutionary strata (blue, purple, orange and black strata;** [43]).

*P* values <0.05 are in bold. NA: not applicable, as the young strata only had one DE gene, statistical comparisons could not be performed for this compartment.

**S9 Table. Wilcoxon rank-sum test statistics for comparisons of unoriented differences in overall GC content (GC0), and third codon position (GC3) between alleles of differentially expressed (DE) versus non-differentially expressed (non-DE) genes within genomic compartments, including autosomes pseudo-autosomal regions (PARs), young evolutionary strata (previously identified red and green strata;** [43]**) and old evolutionary strata (blue, purple, orange and black strata;** [43]).

(A) Overall GC content. (B) Third codon position GC content. *P* values <0.05 are in bold. NA: not applicable, as the young strata only had one DE gene, statistical comparisons could not be performed for this compartment.

**S1 Fig. Heatmap showing differentially expressed genes between haploid a_1_ culture and haploid a_2_ of *Microbotryum lychnidis-dioicae* cultures under low nutrient condition.**

Each column shows a replicated sample for each haploid cell culture. Z-score denotes the relative gene expression level, and cluster shows the similar gene expression profiles.

**S2 Fig. Interaction plots for pairs of predictor variables in overall GLM of differential gene expression between mating types of *Microbotryum lychnidis-dioicae*.**

Y-axes are GLM-predicted response values of differential expression ratio between allele pairs in a_1_ and a_2_ haploid genomes, and x-axes are allele differences between allele pairs in a_1_ and a_2_ haploid genomes in predicted protein length as the predictor variable binned into levels of interacting categorical predictor variable (i.e. panel **A**, genomic compartment) or other interacting continuous predictor variables (i.e. panels **B-D**; the lowest bin being no differences between alleles, and low and high bins being split at the median value among genes with non-zero differences between alleles). (**A**) Interaction plot between protein length differences and genomic compartment. Genomic compartments include autosomes, pseudo-autosomal regions (PARs), young evolutionary strata (previously identified red and green strata; [43]) and old evolutionary strata (blue, purple, orange and black strata; [43]). (**B**) Interaction plots between protein length differences and differences in transposable elements (TEs) insertions. (**C**) Interaction plots between protein length differences and non-synonymous mutation (*dN*) rate differences. (**D**) Interaction plots between protein length differences and GC content differences.

**S3 Fig. Boxplot of synonymous mutation rate (*dS*) for differentially expressed (DE) and non-differentially expressed genes (Non-DE) of *Microbotryum lychnidis-dioicae*.**

Wilcoxon rank sum tests for comparisons of mean non-synonymous mutation rate (*dN*) of differentially expressed genes (DE) versus non-differentially expressed genes (non-DE) within genomic compartments: ‘***’: *P* < 0.001, other comparisons were not significant. Genomic compartments include autosomes, pseudo-autosomal regions (PARs), young evolutionary strata (previously identified red and green strata; [43]) and old evolutionary strata (blue, purple, orange and black strata; [43]).

**S4 Fig. Boxplot of differentially (a_1_-biased, a_2_-biased) and non-differentially expressed genes (not-biased) and the sequence divergence between alleles of *Microbotryum lychnidis-dioicae*.**

Wilcoxon rank sum tests for comparisons of genes with higher allele expression in the a_1_ and a_2_ haploid mating type genomes separately to non-differentially expressed genes for the mean non-synonymous mutation rate (*dN*) (**A**), synonymous mutation rate (*dS*) (**B**); NS: not significant, ‘***’: *P* < 0.001, ‘**’: *P* < 0.01, ‘*’: *P* < 0.5, ‘.’: *P* < 0.1, NS: not significant. As *dN* and *dS* of almost all genes in autosome and PAR are zero, and there is only one DE gene on the young strata, so no statistic test can be performed in these regions. Sample size for each genomic compartment is listed either above or inside boxplot accordingly. Genomic compartments include autosomes, pseudo-autosomal regions (PARs), young evolutionary strata (previously identified red and green strata; [43]) and old evolutionary strata (blue, purple, orange and black strata; [43]).

**S5 Fig. Boxplot of differentially expressed (DE) and non-differentially expressed genes (non-DE) and gene evolutionary rate *dN/dS* of *Microbotryum lychnidis-dioicae*.**

Wilcoxon rank sum tests for comparisons of evolutionary rate *dN*/*dS* of differentially expressed genes (DE) versus non-differentially expressed genes (non-DE) within genomic compartments; NS: not significant. As *dN*/*dS* of almost all genes in autosome and PAR are zero, and there is only one DE gene on the young strata, so no statistic test can be performed in these regions. Genomic compartments include autosomes, pseudo-autosomal regions (PARs), young evolutionary strata (previously identified red and green strata; [43]) and old evolutionary strata (blue, purple, orange and black strata; [43]).

**S6 Fig. Boxplot of sequence divergence, non-synonymous mutation rate *dN* (A), synonymous mutation rate *dS* (B), between *Microbotryum lychinidis-dioicae* and *M. lagerheimii*.**

Alleles of differentially expressed genes with hypothesized higher (red - lower expressed allele) and lower (blue - higher expressed allele) substitution rates and hypothesized equal (grey; non-differentially expressed genes) mutation rates were pooled from a_1_ and a_2_ genomes and assessed for divergences from orthologs in *M. lagerheimii*. Genomic compartments include autosomes, pseudo-autosomal regions (PARs), young evolutionary strata (previously identified red and green strata; [43]) and old evolutionary strata (blue, purple, orange and black strata; [43]).

**S7 Fig. Dotplot of oriented differences of transposable element (TE) insertions and differential gene expression between alleles of *Microbotryum lychnidis-dioicae*.**

TE insertions are shown for sliding-window intervals from upstream to downstream of genes, were differences between alleles were calculated as the TE number for the allele with lower expression minus the TE number for the higher expressed allele; a positive value thus represented an excess of TEs in the lower expressed allele. Sliding window intervals are shown as **A**: upstream 20kb to 10kb, **B**: upstream 15kb to 5kb, **C**: upstream 10kb to gene, **D**: upstream 5kb to downstream 5kb, **E**: gene to downstream 10kb, **F**: downstream 5kb to 15kb, and **G**: downstream 10kb to 20kb.

**S8 Fig. Average indel numbers and proportions of genes with different stop codon positions between alleles of differentially expressed genes of *Microbotryum lychnidis-dioicae*.**

Among genes having alleles with different predicted protein lengths, boxplot of average indel numbers for both differentially expressed (DE) and non-DE genes across various genomic compartments (**A**), and barplot for proportions of genes with different stop codon positions for both DE and non-DE genes across genomic compartments (**B**). **: P < 0.01, *: P < 0.05, ‘.’: P < 0.1, NS: not significant. Genomic compartments include autosomes, pseudo-autosomal regions (PARs), young evolutionary strata (previously identified red and green strata; [43]) and old evolutionary strata (blue, purple, orange and black strata; [43]).

**S9 Fig. Comparisons of differentially expressed (DE) and non-differentially expressed (non-DE) genes between mating types of *Microbotryum lychnidis-dioicae* for differences between alleles in third codon position GC content (GC3) within genomic compartments.**

Wilcoxon rank sum test statistics for comparisons of unoriented differences in GC content of third codon position (GC3) between alleles of differentially expressed (DE) versus non-differentially expressed (non-DE) genes within genomic compartments; ***: *P* < 0.001, NS: non significant. Genomic compartments include autosomes, pseudo-autosomal regions (PARs), young evolutionary strata (previously identified red and green strata; [43]) and old evolutionary strata (blue, purple, orange and black strata; [43]). The notation “*a*” indicates that the young evolutionary strata contained only one DE gene, precluding comparisons to non-DE genes within this compartment.

**S10 Fig. Comparison of proportion (A) and number (B) of differentially expressed (DE) genes detected between a_1_ and a_2_ haploid mating type genomes of *Microbotryum lychnidis-dioicae*.**

Differentially expressed (DE) genes on mating-type chromosome (MAT) chromosomes and autosomes (auto), at various percentage protein sequence identities used as threshold for identification of alleles for genes in a_1_ and a_2_ haploid genomes. **: *P* < 0.01 using Chi-square test. All other comparisons of DE genes on mating-type chromosome and autosomes are not significant.

**S11 Fig. Multidimensional scaling (MDS) plot of RNAseq libraries of *Microbotryum lychnidis-dioicae*.**

Water denotes water agar (i.e. low nutrients) culture condition.

**S12 Fig. The ratio index of coding sequence and protein sequence of *Microbotryum lychnidis-dioicae*.**

The ratio of predicted coding sequence divided by the predicted protein sequence multiplied by three for a_1_ and a_2_ alleles, among four genomic compartments.

## Data access

We used published gene expression data to investigate the association of sequence degeneration and differential gene expression in *Microbotryum lychnidis-dioicae* ([42,45], https://trace.ncbi.nlm.nih.gov/Traces/study/?acc=+PRJNA246470&go=go). We used published genome assembly, gene predictions and assignations to genomic compartments [43,44]. We also used published transposable elements identification in *M. lychnidis-dioicae* [87]. All relevant scripts and data files to perform these analyses are deposited in Zenodo and Github (https://github.com/Wen-Juan/Differential_expression_associateswith_degeneration_Microbotryum_fungus), which will be released immediately upon manuscript acceptance.

## Acknowledgements

We thank Darren J. Parker for assistance on python scripting and valuable discussions of the results, and Ricardo C. Rodríguez de la Vega for insightful comments. The computations were performed at the Vital-IT (http://www.vital-it.ch) Center for high-performance computing of the SIB Swiss Institute of Bioinformatics. This work was supported by the NIH grant R15GM119092 to M. E. H., and the Louis D. Foundation award to T. G.

